# DCBLD1 modulates angiogenesis by regulating of the VEGFR-2 endocytosis in endothelial cells

**DOI:** 10.1101/2023.04.20.537746

**Authors:** Qi Feng, Lingling Guo, Chao Yu, Xiaoning Liu, Yanling Lin, Chenyang Li, Wenjun Zhang, Yanhong Zong, Weiwei Yang, Yuehua Ma, Runtao Wang, Lijing Li, Yunli Pei, Huifang Wang, Demin Liu, Mei Han, Honglin Niu, Lei Nie

## Abstract

**Objective:** Unwanted angiogenesis is involved in the progression of various malignant tumors and cardiovascular diseases, and the factors that regulate angiogenesis are potential therapeutic targets. We tested the hypothesis that DCBLD1 (Discoidin, CUB, and LCCL domain-containing protein 1) is a co-receptor of VEGFR-2 and modulates angiogenesis in endothelial cells(ECs).

**Approach and Results:** A carotid artery ligation model and retinal angiogenesis assay were used to study angiogenesis using globe knockout or EC-specific conditional DCBLD1 knockout mice *in vivo*. Immunoblotting, immunofluorescence staining, plasma-membrane subfraction isolation, Co-immunoprecipitation and mass-spectrum assay were performed to clarify the molecular mechanisms.Loss of DCBLD1 impaired VEGF response and inhibited VEGF-induced EC proliferation and migration. DCBLD1 deletion interfered with adult and developmental angiogenesis. Mechanistically, DCBLD1 bound to VEGFR-2 and regulated the formation of VEGFR-2 complex with negative regulators: protein tyrosine phosphatases, E3 ubiquitin ligases(Nedd4 and c-Cbl), and also DCBLD1 knockdown promoted lysosome-mediated VEGFR-2 degradation in ECs.

**Conclusions:** These findings demonstrated the essential role of endothelial DCBLD1 in regulating VEGF signaling and provided evidence that DCBLD1 promotes VEGF-induced angiogenesis by limiting the dephosphorylation, ubiquitination, and lysosome degradation after VEGFR-2 endocytosis. We proposed that endothelial DCBLD1 is a potential therapeutic target for ischemic cardiovascular diseases by the modulation of angiogenesis through regulating of the VEGFR-2 endocytosis.

## Introduction

Angiogenesis is the formation of new blood vessels from pre-existing blood vessels^1–3^. This biological process begins with the sprouting of endothelial cells (ECs) in preexisting capillaries, followed by EC migration, proliferation, and tube formation^4^. Pathological angiogenesis occurs in various forms of malignant tumors and cardiovascular diseases^5^.

VEGF is a key regulator of physiological angiogenesis during embryo-genesis, bone growth and reproductive function, and pathological angiogenesis related to tumors, intraocular new vascular, and other diseases^6^. VEGF-A is the main soluble agent that drives angiogenesis, which activates tyrosine kinase receptor family VEGFRs^7^. Although VEGF-A binds to VEGFR1 and VEGFR2, the signaling is mainly mediated by VEGFR2, resulting in vascular permeability and EC proliferation, survival, and migration^8^. These pathophysiological changes involve the initiation of multiple signal transduction pathways, such as PI3K-Akt, p44/42MAPK, and calcium signaling, to transduce the activation of VEGF signaling^9^.

The discoidin, CUB, and LCCL domain-containing (DCBLD) receptor family comprises the type-I transmembrane proteins, DCBLD1 and DCBLD2 (also known as ESDN or CLCP1). The DCBLD receptor family plays an important role in development and cancer^10^. DCBLD1 is a gene encoding unknown membrane proteins, which are over-expressed in lung adenocarcinoma. Additionally, it is reported to be a prognostic factor for overall survival of non-small cell lung carcinoma (NSCLC) and breast cancer^11,12^. The expression of DCBLD1 can be inhibited by creating YY1 binding sites, which reduce the risk of lung adenocarcinoma^12^. In this regard, non-receptor tyrosine kinases, FYN and ABL, can phosphorylate tyrosine residues in the YXXP motif in the intracellular domain of the members of the DCBLD family. The phosphorylation results in the recruitment of signaling proteins with Src homologous 2 (SH2) domain (CRK) and CRK-like (CRKL) of the kinase CT10 regulator interacting with the new scaffold activated receptors, DCBLD1 and DCBLD2^13,14^. The DCBLD receptor family is highly conserved across vertebrates and contain similar domain structures. Previous results have demonstrated that DCBLD2 alters VEGF function and angiogenesis in endothelial cells^15^. We hypothesized that the structural similarity between DCBLD1 and DCBLD2 increases the possibility that it may regulate VEGF signaling activities on ECs.

In this study, we demonstrated that DCBLD1 regulates VEGF-induced EC proliferation, migration, *in vivo* angiogenesis during development, and adult angiogenesis in mice. DCBLD1 accomplished these roles by regulating the homeostasis of VEGFR-2 after endocytosis or the formation of VEGFR-2-protein tyrosine phosphatase/E3 ubiquitin ligase complex. These findings revealed that DCBLD1 mediated its function by regulating the VEGF signal transduction in ECs, suggesting that DCBLD1 could be a potential target for regulating angiogenesis.

## Results

### DCBLD1 is expressed in ECs and regulates the proliferation, migration, and tube formation of ECs in response to VEGF

We confirmed the expression of DCBLD1 in human umbilical vein endothelial cells (HUVECs) or mouse pulmonary microvascular endothelial cells (mPMVECs) by Western blot (Fig. 1A-D) and immunofluorescence staining (Fig. 1E). DCBLD1 was localized on the cell membrane in HUVECs (Fig. 1E). The Western blot showed that DCBLD1 protein levels were significantly higher in proliferating non-confluence HUVECs than in growth-arrested confluence HUVECs (Fig. 1C, D).

**Figure 1.**
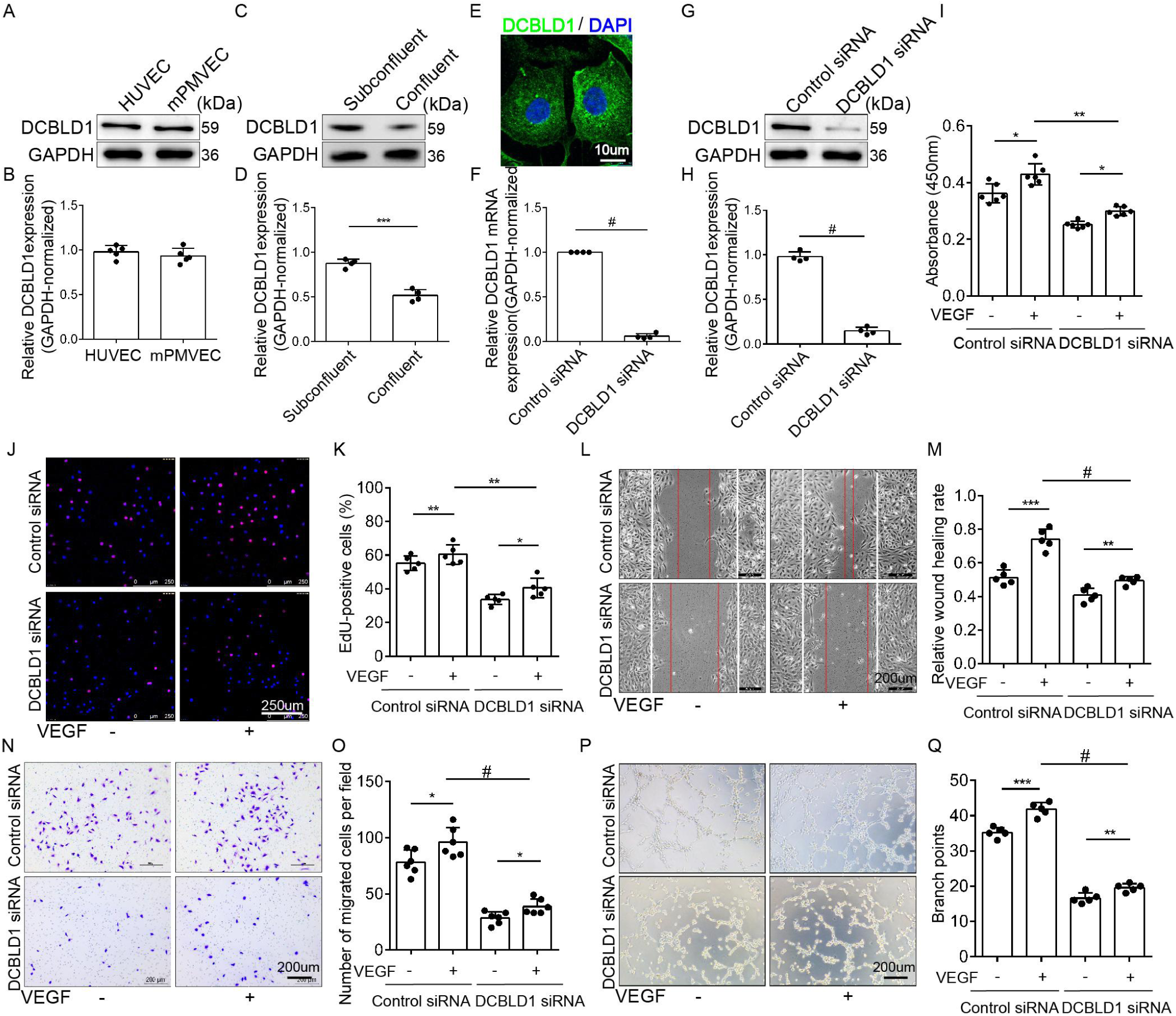
DCBLD1 modulation of VEGF-induced endothelial cell proliferation, migration, and tube formation. **(A, B)** The expression of DCBLD1 protein in HUVECs and mPMVECs was analyzed by Western blot (*n*=5*, p*=0.3936). **(C, D)** Western blot was used to detect the expression of GAPDH-normalized DCBLD1 protein in HUVECs under sub-confluent and confluent conditions (*n*=4, \*\*\**P*<0.001). **(E)** Microscopic images of DCBLD1 immunofluorescence staining (green) in HUVECs. The nucleus was labeled blue with DAPI. Bar=10 μm. **(F-H)** Real-time RT-PCR (F, *n*=4, ^#^*P*<0.0001) and western blot (G and H, *n*=4, ^#^*P*<0.0001) were used to detect the expression of GAPDH-normalized DCBLD1 protein and mRNA in HUVECs after DCBLD1 siRNA knockdown. **(I)** HUVECs were subjected to siRNA-mediated DCBLD1-knockdown, and VEGF-induced EC growth was detected by BrdU. The absorbance at 450nm (OD450) was detected by microplate reader to detect the content of BrdU incorporated into DNA, so as to judge the proliferation ability of cells (*n*=6, \**P*<0.05, \*\**P*<0.01). **(J, K)** VEGF-induced EC proliferation was monitored by EdU assay after siRNA-mediated DCBLD1 knockdown (*n*=5, \**P*<0.05, \*\**P*<0.01). Bar=250 μm.**(L, M)** HUVECs were subjected to siRNA-mediated DCBLD1-knockdown, and VEGF-induced EC migration was analyzed by scratch assay (*n*=5, \*\**P*<0.01,\*\*\**P*<0.001,^#^*P*<0.0001). Bar=200 μm. **(N, O)** VEGF-induced EC migration was monitored by Transwell assay after siRNA-mediated DCBLD1 knockdown (*n*=6, \**P*<0.05, ^#^*P*<0.0001). Bar=200 μm. **(P, Q)** HUVECs were subjected to siRNA-mediated DCBLD1-knockdown, and VEGF-induced EC angiogenesis was analyzed by tubule formation experiment (*n*=5, \*\**P*<0.01,\*\*\**P*<0.001,^#^*P*<0.0001) Bar= 200 μm. The values are represented by mean±SEM. Statistical analyses were performed using two-way ANOVA, followed by Tukey’s multiple comparison test, compared to the control at the same time point.

Based on the structural similarity between DCBLD1 and DCBLD2, we investigated the potential role of DCBLD1 in regulating VEGF response in ECs. We investigated the effects of DCBLD1 on EC proliferation, migration, and tube formation by knocking-down the expression of DCBLD1. Quantitative RT-PCR (Fig. 1F) and Western blot (Fig. 1G, H) were used to evaluate the efficiency of siRNA transfection on DCBLD1 expression in HUVECs. The BrdU (Fig. 1I) and EdU (Fig. 1J, K) experiments showed that DCBLD1 knockdown decreased the proliferation of HUVECs induced by VEGF. The scratch experiment showed that DCBLD1 knockdown reduced the migration ability of HUVECs (Fig. 1L, M). The transwell chamber migration experiment showed that DCBLD1 knockdown decreased the transmembrane migration ability of HUVECs (Fig. 1N, O). The tube formation assay is a proxy for angiogenetic activities of ECs *in vitro*^15^. The results showed that DCBLD1 knockdown reduced the tube-forming ability of HUVECs in response to VEGF (Fig. 1P, Q). Together, these data support the hypothesis that DCBLD1 plays an essential role in VEGF response regulation in HUVECs.

### DCBLD1 gene knockdown or deletion inhibits VEGF signaling in ECs

To clarify the exact mechanism of the effect of DCBLD1 on VEGF response in ECs, we studied the effect of downregulation or knockout of DCBLD1 on VEGF signal transduction events. Western blot analysis showed that VEGF-induced activation of VEGFR2, Akt, MAPK p44/42, and p38 was reduced in DCBLD1 knockdown HUVECs (Fig. 2A-F) or DCBLD1 knockout mPMVECs (*Dcbld1*^-/-^ mPMVECs) (Fig. 2G-L). Together, these data indicated that the lack of DCBLD1 impaired VEGF signal transduction in ECs.

**Figure 2.**
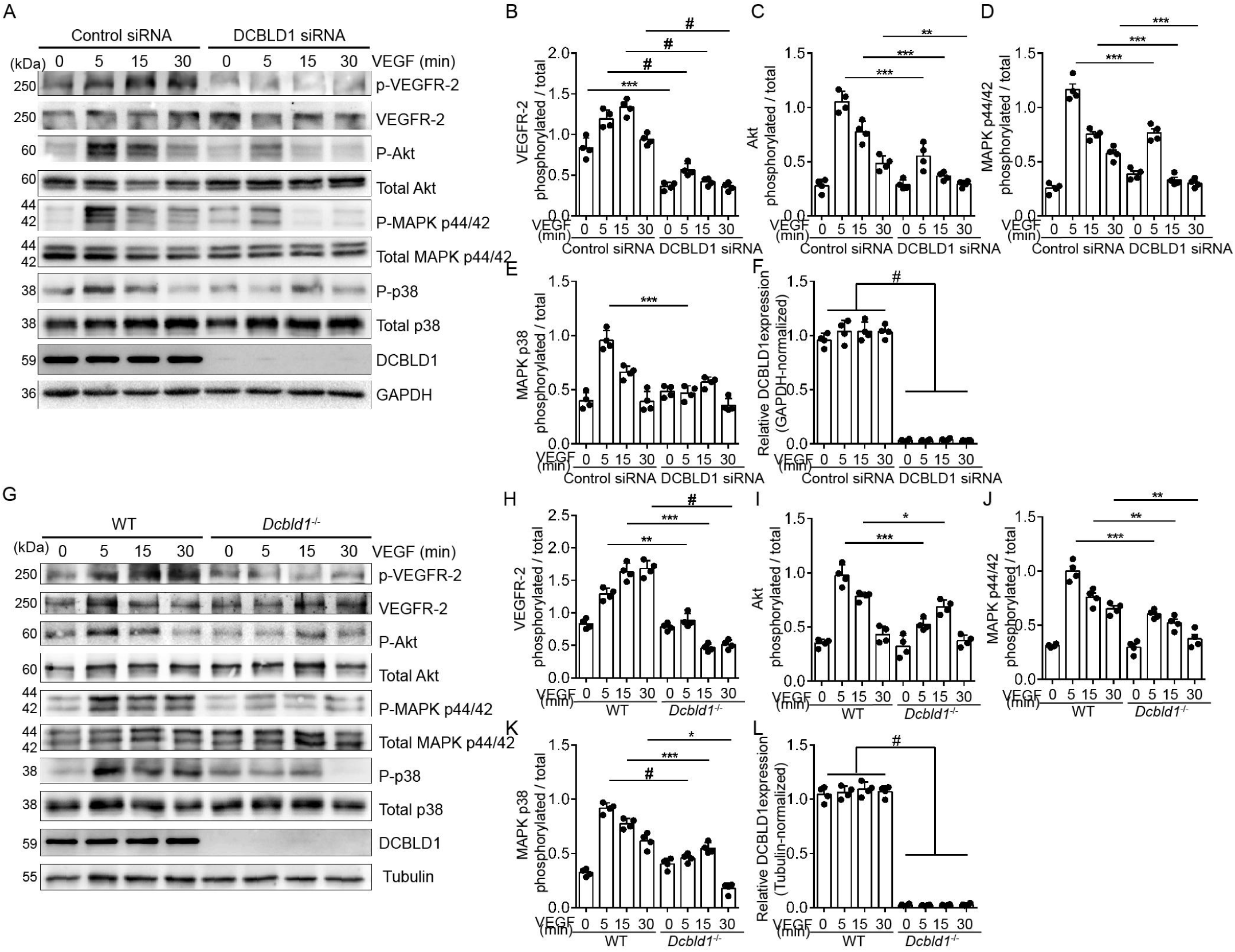
DCBLD1 Deletion inhibited VEGF signaling pathway in ECs. **(A-F)** Western blot of proteins implicated in VEGF signaling in HUVECs. Representative Western blot and quantification of p-VEGFR-2, MAPK P-p44/42 Thr^202^/Tyr^204^, P-Akt Ser^473^ and MAPK P-p38 in HUVECs treated with VEGF (10 ng/mL) at different time points, following transient transfection with DCBLD1 siRNA. GAPDH was used as a loading control (*n=*4/group.\*\*\**p<*0.001, ^#^*p<*0.0001).**(G-L)** Western blot of proteins in relation to VEGF signaling in mPMVECs isolated from WT and *Dcbld1^-/-^*mice. Representative Western blot and quantification of p-VEGFR-2, MAPK P-p44/42 Thr^202^/Tyr^204^, P-Akt Ser^473^, and MAPK P-p38 in ECs treated with VEGF (10 ng/mL) at different time points. Tubulin was used as a loading control (*n=*4/group. \**p<*0.05, \*\**p<*0.01, \*\*\**p<*0.001, ^#^*p<*0.0001). Data points indicate mean±SEM. Statistical analyses were performed using two-way ANOVA, followed by Tukey’s multiple comparisons test. The values are represented by mean±SEM. Statistical analysis: Two-way ANOVA followed by Tukey’s multiple comparison test, compared to the control at the same time point.

### DCBLD1 deletion promoted intimal hyperplasia after carotid artery ligation

From the experiments described above, we proved that the loss of DCBLD1 impaired EC function. Studies have shown that EC dysfunction significantly increases intimal hyperplasia^16^. To evaluate the effect of DCBLD1 on intimal hyperplasia, we developed DCBLD1 knockout (*Dcbld1*^-/-^) mouse models and endothelial-specific DCBLD1 knockout (Tek-*Cre*^+/-^*Dcbld1*^flox/flox^) mouse models. We used specific primers for PCR and agarose gel electrophoresis of mouse genomic DNA. The results showed that the *Dcbld1*^-/-^ mice had no bands at 225 bp. The *Dcbld1*^flox/flox^ mice showed a clear product of *lox*P at 468 bp/484 bp, and the *Dcbld1*^flox/+^ mice showed a *lox*P heterozygous product at 468 bp/484 bp and 356 bp/377 bp. The Tek-*Cre*^+/−^*Dcbld1*^flox/flox^ mice showed a clear product of *lox*P at 468 bp/484 bp and a positive product of *Cre* sequence at 100 bp molecular weight marker (Fig. 3A). These findings indicate that the genotypic identification of the mice to be used in subsequent experiments was correct, and the DCBLD1 knockout mice and endothelial-specific DCBLD1 knockout mice were successfully developed.

**Figure 3.**
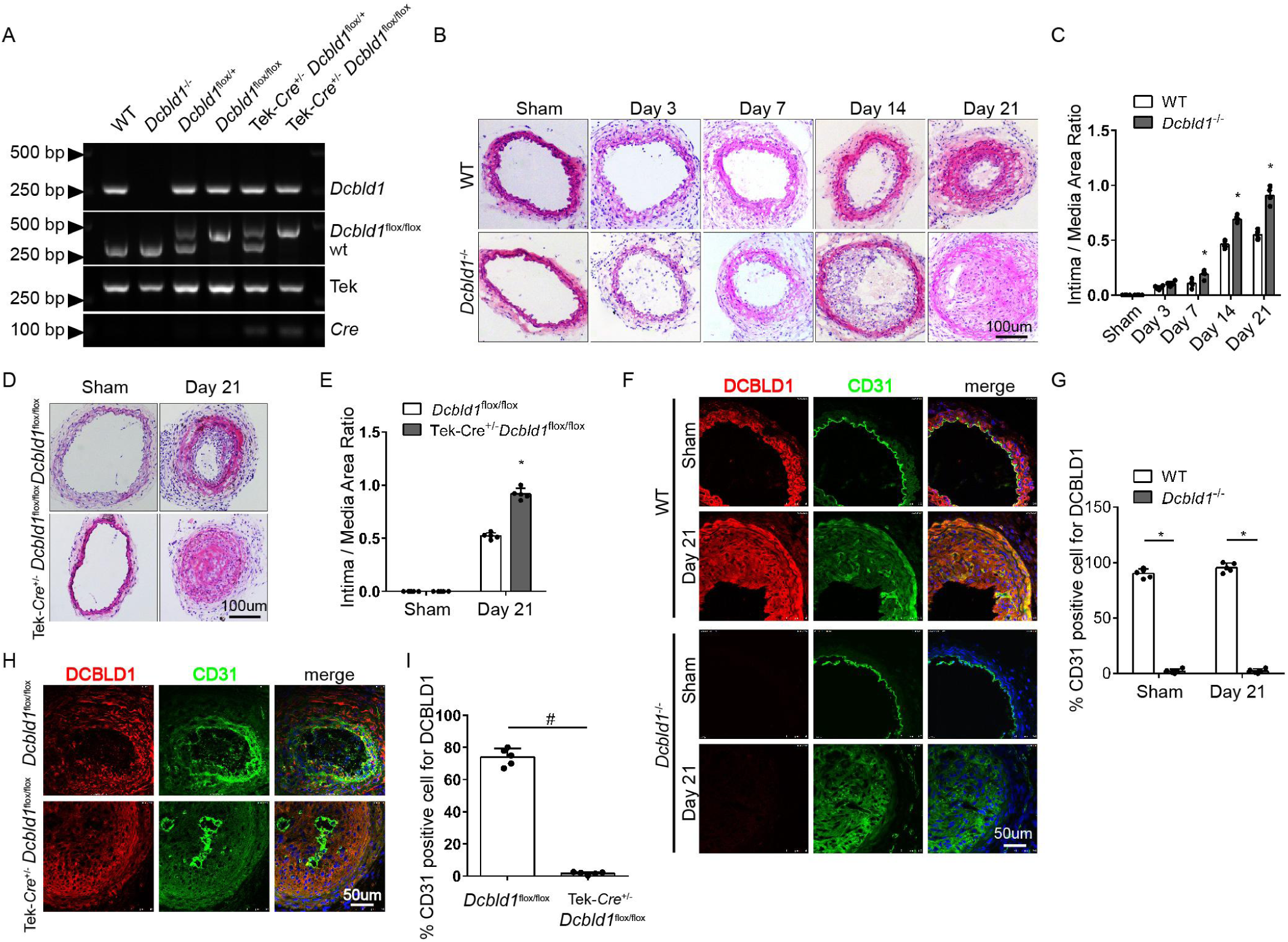
DCBLD1 deletion promoted intimal hyperplasia after carotid artery ligation. **(A)** The genotype PCR products were subjected to agarose gel electrophoresis. **(B-E)** The quantitative results of H&E staining on the representative sections and the ratio of intima-to-membrane of WT and *Dcbld1^-/-^* mice (B, C) or Tek-*Cre*^+/−^*Dcbld1*^flox/flox^ mice **(D, E)** at different time points after carotid artery ligation were summarized. Each point represents a single mouse; the data points indicate mean±SEM, Bar=100 μm, *n=*5/group, \**p<*0.05. Statistical analysis: Two-way ANOVA followed by Tukey’s multiple comparison test. **(F-I)** Immunofluorescence staining of representative sections of common carotid artery in WT and *Dcbld1*^-/-^ mice **(F, G)** or Tek-*Cre*^+/−^*Dcbld1*^flox/flox^ mice **(H, I)** at 21 days after carotid artery ligation. Color overlay represents the expression of DCBLD1 in endothelial cells. Bar=50 μm. *n=*5/group, \**p<*0.05, ^#^*p<*0.0001. The values are represented by mean±SEM. Statistical analysis were performed using two-way ANOVA, followed by Tukey’s multiple comparison test, compared to the control at the same time point.

We examined the role of DCBLD1 in mouse intimal hyperplasia using a common carotid artery ligation model. The effect of DCBLD1 expression on intimal hyperplasia after carotid artery ligation was determined by H&E staining at different time points. The results showed that compared with WT mice, *Dcbld1*^-/-^ mice had significantly increased intimal thickening at the same time interval, which reached the peak 21 days after ligation (Fig. 3B, C). The effect of DCBLD1 expression in ECs on intimal hyperplasia was measured by EC-specific DCBLD1 knockout mice. The results showed that compared with *Dcbld1*^flox/flox^ mice, Tek-*Cre*^+/−^*Dcbld1*^flox/flox^ mice had significantly increased intimal thickening on Day 21 (Fig. 3D,E).

The distribution of DCBLD1 expression was analyzed by immunofluorescence staining, and the effect of DCBLD1 on intimal hyperplasia was verified. From the vascular tissues of WT mice, DCBLD1 was expressed in the intima and middle and outer membranes of the blood vessel; whereas the expression of DCBLD1 in vascular tissues of *Dcbld1^-/-^*mice was low, indicating that *Dcbld1^-/-^* mice were successfully developed (Fig. 3F, G). At 21 days after ligation, the degree of intimal hyperplasia in *Dcbld1^-/-^* mice was significantly higher than that in WT mice, indicating that DCBLD1 knockout promotes intimal hyperplasia after carotid artery ligation (Fig. 3F). Immunofluorescence staining of carotid artery in Tek-*Cre*^+/−^*Dcbld1*^flox/flox^ mice revealed that the expression of DCBLD1 was significantly downregulated in the CD31 positive region of EC-specific marker, indicating that Tek-*Cre*^+/−^*Dcbld1*^flox/flox^ mice were successfully developed (Fig. 3H, I). At 21 days after ligation, the degree of carotid intimal hyperplasia in Tek-*Cre*^+/−^*Dcbld1*^flox/flox^ mice was significantly higher than that in control mice (Fig. 3H).

### DCBLD1 deletion inhibits angiogenesis in adult and developing mice

To investigate the effect of DCBLD1 on angiogenesis *in vivo*, we used 6–8 weeks old male mice to establish hind limb ischemia (HLI) models induced by femoral artery ligation and resection. Thereafter, we observed the recovery of lower limb blood flow by laser Doppler perfusion imaging to determine the effect of angiogenesis in mice, as described in literature^17,18^. The results showed that the hind limb blood flow recovery of WT mice increased significantly on the 7th day after ischemia and approached the contralateral hind limb blood flow recovery on the 14th day (Fig. 4A, B). However, the blood flow recovery in *Dcbld1*^-/-^ mice was significantly delayed, compared to that in WT mice. Immunofluorescence analysis showed that angiogenetic markers, CD31 (Fig.4E, F) and NG2 (Fig.4G, H), were significantly decreased in the gastrocnemius of *Dcbld1*^-/-^ mice on Day 14 after HLI, compared with that of WT mice. These results showed that the deletion of DCBLD1 inhibited the angiogenesis of adult mice in the HLI model.

**Figure 4.**
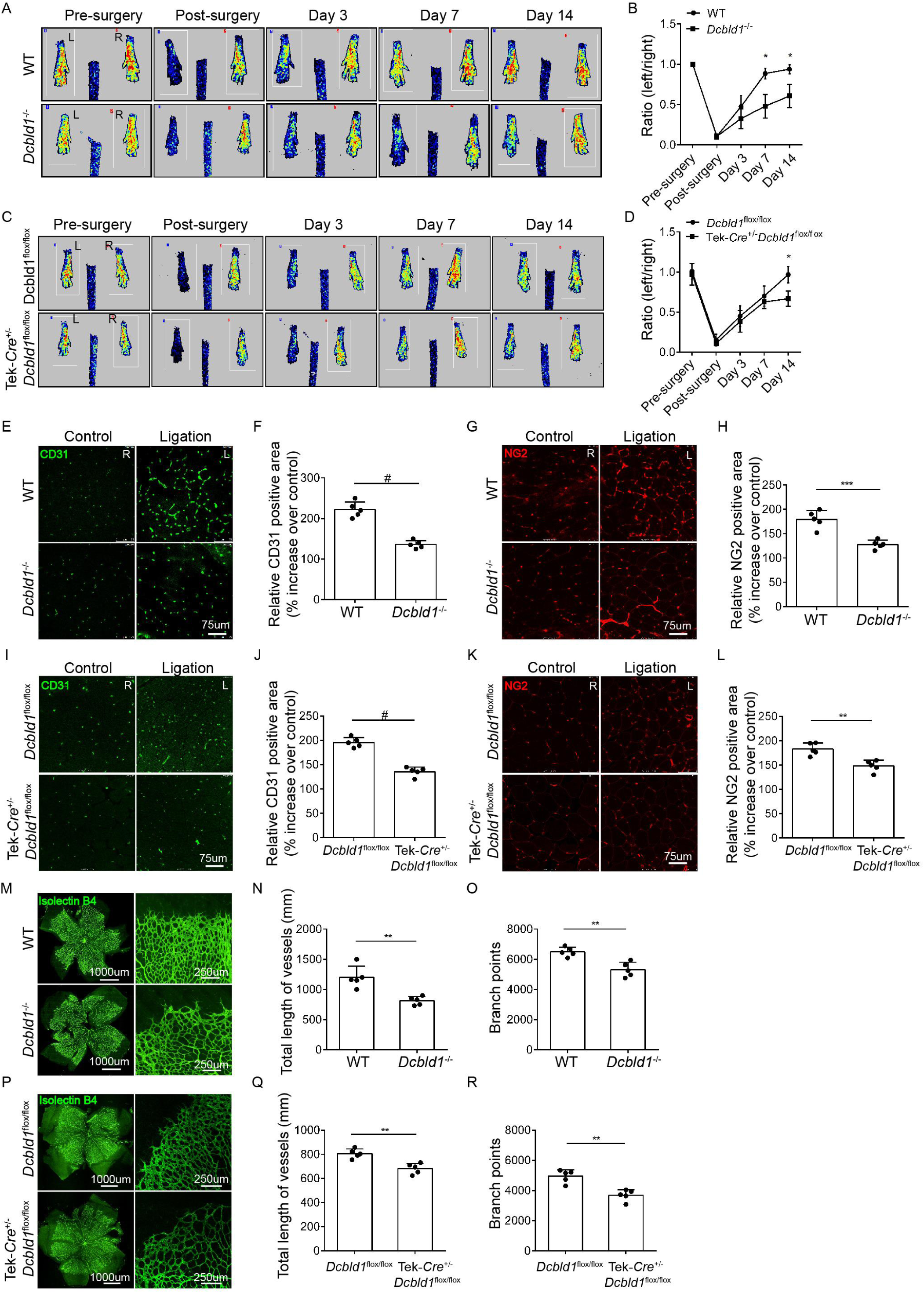
DCBLD1 deletion inhibits angiogenesis in adult and developing mice. **(A, B)** Representative examples of laser Doppler images before and after ligation of left femoral artery in WT and *Dcbld1^-/-^* mice and quantification of hind limb blood flow. *n=*5, \**p<*0.05. **(C, D)** Representative examples of laser Doppler images before and after ligation of left femoral artery in Tek-*Cre*^+/−^*Dcbld1*^flox/flox^ mice and quantification of hind limb blood flow. *n=*5, \**p<*0.05. **(E, F)** Representative examples and quantification of CD31 immunostaining in gastrocnemius of WT or *Dcbld1^-/-^*mice 14 days after femoral artery ligation, Bar=75 μm, *n=*5, ^#^*p<*0.0001. **(G, H)** Representative examples and quantification of NG2 immunostaining in gastrocnemius of WT or *Dcbld1^-/-^* mice 14 days after femoral artery ligation, Bar=75 μm, *n=*5, \*\*\**p<*0.001. **(I, J)** Representative examples and quantification of CD31 immunostaining in gastrocnemius of *Dcbld1*^flox/flox^ or Tek-*Cre*^+/−^*Dcbld1*^flox/flox^ mice 14 days after femoral artery ligation, Bar=75 μm, *n=*5, ^#^*p<*0.0001. **(K, L)** Representative examples and quantification of NG2 immunostaining in gastrocnemius of *Dcbld1*^flox/flox^ or Tek-*Cre*^+/−^*Dcbld1*^flox/flox^ mice 14 days after femoral artery ligation, Bar=75 μm, *n=*5, \*\**p*<0.01. **(M-O)** The representative examples of Isolectin B4 staining and retinal vascular quantification in WT and *Dcbld1^-/-^* mice on the fifth day after birth. Bar: 1000 μm upper panel, 250 μm lower panel. *n=*5, \*\**P<*0.01. **(P-R)** The representative examples of Isolectin B4 staining and retinal vascular quantification in *Dcbld1*^flox/flox^ and Tek-*Cre*^+/−^*Dcbld1*^flox/flox^ mice on the fifth day after birth. Bar: 1000 μm upper panel, 250 μm lower panel. *n=*5, \*\**p<*0.01.The values are represented by mean±SEM. Statistical analyses were performed using two-way ANOVA, followed by Tukey’s multiple comparison test, compared to the control at the same time point.

The development of retinal vessels in mice began from birth and ended at the age of 1 week. The endothelial cell marker, Isolectin B4, was used for immunofluorescence staining on the retinal vessels of 5-day old mice. The total length of retinal vessels and number of vascular branches were measured to indirectly reflect the developmental angiogenetic ability of mice. The results showed that, compared with WT mice, *Dcbld1*^-/-^ mice had significantly decreased total length and number of branches of retinal vessels (Fig. 4M-O). This finding suggested that the deletion of DCBLD1 significantly inhibited the developmental angiogenetic ability of young mice.

### DCBLD1 deletion in endothelial cells inhibits angiogenesis in mice

Furthermore, we used Tek-*Cre*^+/−^*Dcbld1*^flox/flox^ mice to verify the effect of DCBLD1 deficiency in endothelial cells on ischemia-induced angiogenesis. The results showed that, compared with *Dcbld1*^flox/flox^ mice, Tek-*Cre*^+/−^*Dcbld1*^flox/flox^ mice had significantly impaired perfusion recovery of HLI (Fig. 4C, D). Immunofluorescence staining of the gastrocnemius showed that, the CD31 (Fig. 4I, J) and NG2 (Fig. 4K, L) positive cells of Tek-*Cre*^+/−^*Dcbld1*^flox/flox^ mice were significantly decreased, compared with those of*Dcbld1*^flox/flox^ mice on the 14th day of HLI. Isolectin B4 was used to perform immunofluorescence staining on the retinal vessels of 5-day old mice. The results showed that the total length and number of branches of retinal vessels in Tek-*Cre*^+/−^*Dcbld1*^flox/flox^ mice were significantly lower than those in *Dcbld1*^flox/flox^ mice (Fig. 4P-R). The results suggested that DCBLD1 deficiency in ECs significantly inhibited angiogenesis in mice.

### Identification of the VEGFR2 interacting proteins and bioinformatics analyses

Two hundred and thirty immunoprecipitated proteins were successfully identified. Then, the most likely associations between these target proteins and VEGFR2 protein after DCBLD1 knockout were computed by the manual threshold approach and a probabilistic protein-protein interaction prediction algorithm, which identified 48 high-confidence candidates. Supplementary Table 1 shows these proteins were at least 2 fold more or less than 0.5.between DCBLD1 and WT groups. The top-10 enriched in the Gene Ontology (GO) terms of Biological Process (BP), Cellular Component (CC) and Molecular Function (MF) were listed (Supplementary Figure 1). The results of GO showed that target proteins were enriched in several BPs, including endomembrane system organization, plasma membrane orgarization, and transcytosis. Correspondingly, the GO-enriched terms of CCs included membrane raft, coveola, and clathrin-coated endocytic vesicle, For the MF, the majority of the identified proteins were classified into protein tyrosine kinase activity, protein-membrane adaptor activity, and ubiquitin-specific protease binding. Together, the bioinformatic analysis revealed that DCBLD1 involved in membrane receptor endocytosis, adaptor binding, ubiquitination and tyrosine kinase activity.

### DCBLD1 knockdown promoted lysosome-mediated degradation after endocytosis of VEGFR-2 in ECs

Binding of VEGF to VEGFR-2 activates receptor phosphorylation before the receptor enters the cytoplasm by endocytosis^19,20^. The distribution of VEGFR-2 in different components of HUVECs was examined by extracting membrane and cytoplasmic protein fractions. After DCBLD1 knockdown, the level of VEGFR-2 on the cell membrane was significantly decreased, whereas the level of VEGFR-2 in the cytoplasm significantly increased (Fig. 5A-C). These findings indicate that knockdown of DCBLD1 enhances the endocytosis of VEGFR2.

**Figure 5.**
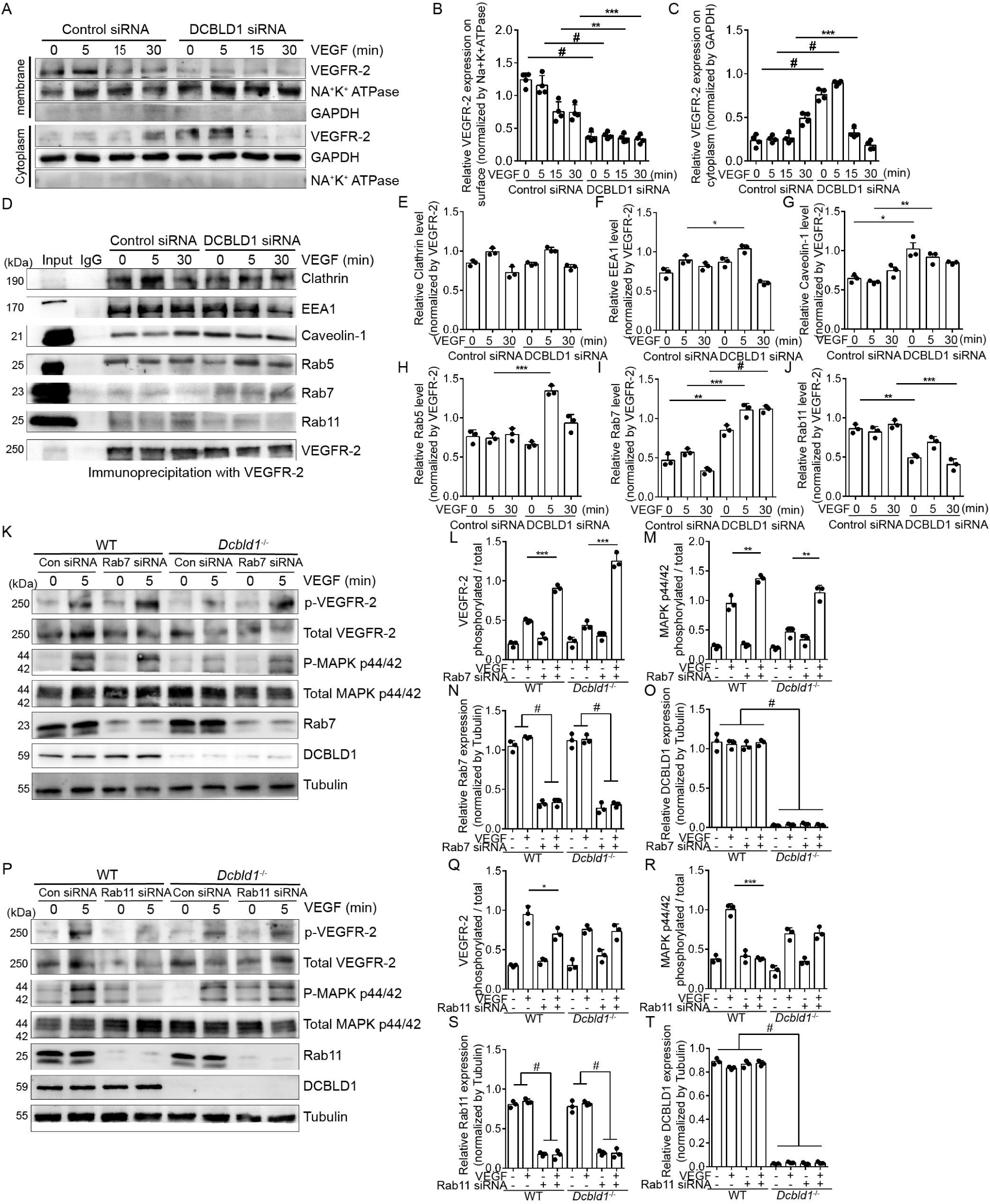
DCBLD1 knockdown accelerated VEGFR-2 endocytosis and lysosomal degradation in ECs. **(A-C)** HUVECs were transiently transfected with DCBLD1 siRNA and then treated with VEGF (10 ng/mL). The distribution of VEGFR-2 between the cytoplasm and membrane components was analyzed at different time points. Na^+^-K^+^ ATPase served as membrane protein loading, and GAPDH as cytoplasmic protein fraction. Western blot showed that DCBLD1 knockdown significantly increased the level of VEGFR-2. *n*=3,\*\**p<*0.01, \*\*\**p<*0.001, ^#^*p<*0.0001. **(D-J)** HUVECs receiving DCBLD1 siRNA were treated with VEGF (10 ng/mL) at different time points. Co-binding analysis of VEGFR-2 with endocytosis marker proteins (clathrin and caveolin-1) and vesicle trafficking-related proteins (EEA1, Rab5, Rab7, and Rab11) in cytoplasm was performed by immunoprecipitation. *n=*3, \**p<*0.05, \*\**p<*0.01, \*\*\**p<*0.001, ^#^*p<*0.0001.**(K-T)** *Dcbld1^-/-^*mPMVEC knockdown of Rab7 (K-O) or Rab11 (P-T), representative Western blot images of VEGF-induced VEGFR-2 and p44/42 Thr^202^/Tyr^204^ MAPK. *n=*3,\**p<*0.05, \*\**p<*0.01, \*\*\**p<*0.001, ^#^*p<*0.0001. The values are represented by mean±SEM. Statistical analyses were performed using two-way ANOVA, followed by Tukey’s multiple comparison test, compared to the control at the same time point.

Recent studies have shown that VEGFR-2 on the cell surface binds to VEGF and enters the cytoplasm mainly by the Clathrin-dependent or Caveolar-dependent endocytosis pathway^21^. After VEGFR-2 entry into cells, the early endosome formed contains the Rab5 marker protein. Rab5 recruits the intracellular adapter protein, APPL, to the endosome. Thereafter, early endosome antigen 1 (EEA1) competitively binds to the same site of Rab5 with APPL. EEA1 gradually replaces the APPL bound to Rab5, which ultimately determines the subcellular localization of endosome^21^. The endosomes or endosomes with Rab11 or Rab4 positive fusion return to the cell membrane and complete the VEGFR-2 recycling process. Alternatively, the marker of endosomes, Rab5, was replaced by Rab7, transported to the multi-vesicular body (MVB), and finally degraded by binding to lysosomes^22–26^. Retention of endosomes in Rab5 vesicles increases the dephosphorylation or ubiquitination of VEGFR-2, which further affects the delocalization of VEGFR-2. In HUVECs, the co-binding status of VEGFR-2 with the endocytosis pathway marker proteins (Clathrin and Caveolin-1), early endocytosis marker proteins (Rab5 and EEA1), recycling endocytosis marker protein (Rab11), and MVB marker protein (Rab7) were analyzed by immunoprecipitation experiments to explore the effect of DCBLD1 on the subcellular localization of VEGFR-2. The co-immunoprecipitation assay showed that knockdown of DCBLD1 accelerated the endocytosis of VEGFR-2 mediated by Caveolin-1. The endocytic VEGFR-2 was markedly transported to MVB and eventually degraded, while a few returned to the cell membrane (Fig. 5D-J). Immunofluorescence labeling revealed that knockdown of DCBLD1 accelerated the binding of VEGFR-2 to Rab7 (Supplemental Fig. 2A, B) and decreased the binding of VEGFR-2 to Rab11 (Supplemental Fig. 2C, D).

Furthermore, through knockdown of Rab7 or Rab11 by siRNA in *Dcbld1*^-/-^ mPMVECs, the effect of Rab7 or Rab11 on the VEGF signaling pathway was observed. The results showed that downregulation of Rab7 reversed the inhibition of VEGFR-2 and p44/42 MAPK phosphorylation induced by DCBLD1 deficiency (Fig 5K-O). Whereas downregulation of Rab11 had minimal effect on VEGF-induced phosphorylation of VEGFR-2 and p44/42 MAPK in *Dcbld1^-/-^*mPMVECs (Fig 5P-T). These results confirmed the previous findings of cellular immune coprecipitation and immunofluorescence, suggesting that the deletion of DCBLD1 promoted Rab7-mediated lysosome degradation after endocytosis of VEGFR-2 in ECs and reduced Rab11-mediated VEGFR-2 recycling to the cell membrane.

### DCBLD1 knockdown promotes dephosphorylation and ubiquitination of VEGFR-2 in ECs

The VEGFR-2 signal is regulated by the binding of VEGFR-2 to many key regulatory proteins, including protein tyrosine phosphatases, such as protein tyrosine phosphatase 1B (PTP1B) and T cell protein tyrosine phosphatase (TCPTP). Additionally, VEGFR-2 can accept ubiquitination modification. Nedd4 and c-Cbl are common E3 ubiquitin ligases of VEGFR-2, which can mediate ubiquitination degradation^27–29^. The inhibition of DCBLD1 downregulation on VEGF signaling and increase of VEGFR-2 endosome binding to Rab5 vesicles prompted us to explore whether the deletion of DCBLD1 would increase the dephosphorylation or ubiquitination of VEGFR-2. In mPMVECs, the co-binding of VEGFR-2 with PTP1B, TCPTP, Nedd4, and c-Cbl was studied in an immunoprecipitation experiment. Western blot showed that compared with WT mPMVECs, *Dcbld1^-/-^*mPMVECs had enhanced binding of VEGFR-2 to phosphatase (Fig. 6A-C) or E3 ubiquitin ligase (Fig. 7A-C) after DCBLD1 deletion. Therefore, we surmised that deletion of DCBLD1 promotes dephosphorylation and ubiquitination of VEGFR-2 in ECs.

**Figure 6.**
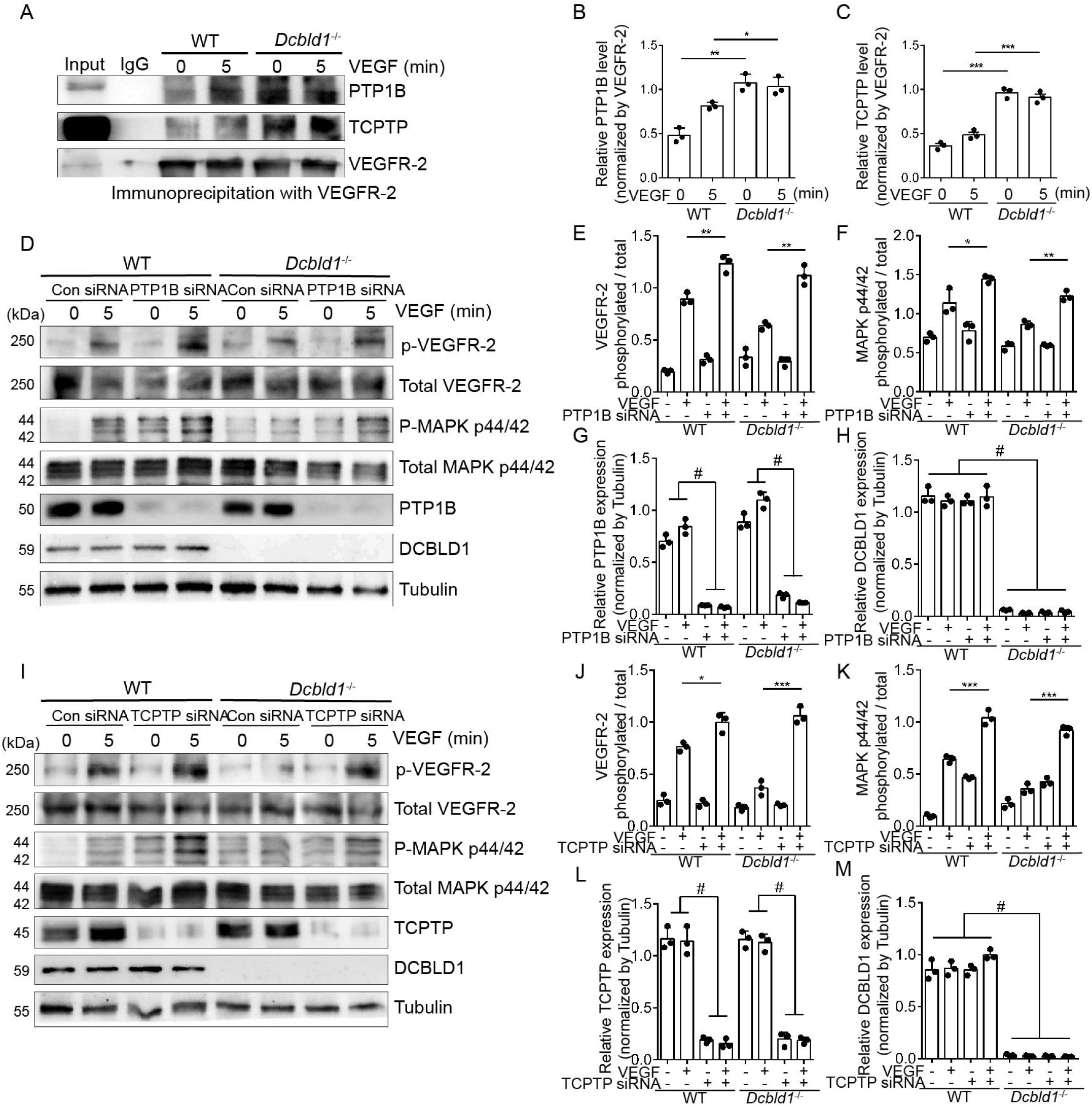
DCBLD1 deletion enhances dephosphorylation of VEGFR-2 and influences VEGF signaling pathway in ECs. **(A-C)** Representative images of Western blot analysis after co-immunoprecipitation of protein tyrosine phosphatase, PTP1B or TCPTP, with VEGFR-2 in WT and *Dcbld1^-/-^*mPMVECs. *n=*3, \**p<*0.05, \*\**p<*0.01, \*\*\**p<*0.001.**(D-M)** Downregulated the expression of PTP1B (D-H) or TCPTP (I-M) proteins in WT and *Dcbld1^-/-^*mPMVECs by siRNA, analyzed the VEGFR-2 and p44/42 Thr^202^/Tyr^204^ MAPK phosphorylation induced by VEGF by Western blot, and performed quantitative analysis. *n=*3, \**p<*0.05, \*\**p<*0.01, \*\*\**p<*0.001, ^#^*p<*0.0001. The values are represented by mean±SEM. Statistical analyses were performed using two-way ANOVA, followed by Tukey’s multiple comparison test, compared to the control at the same time point.

**Figure 7.**
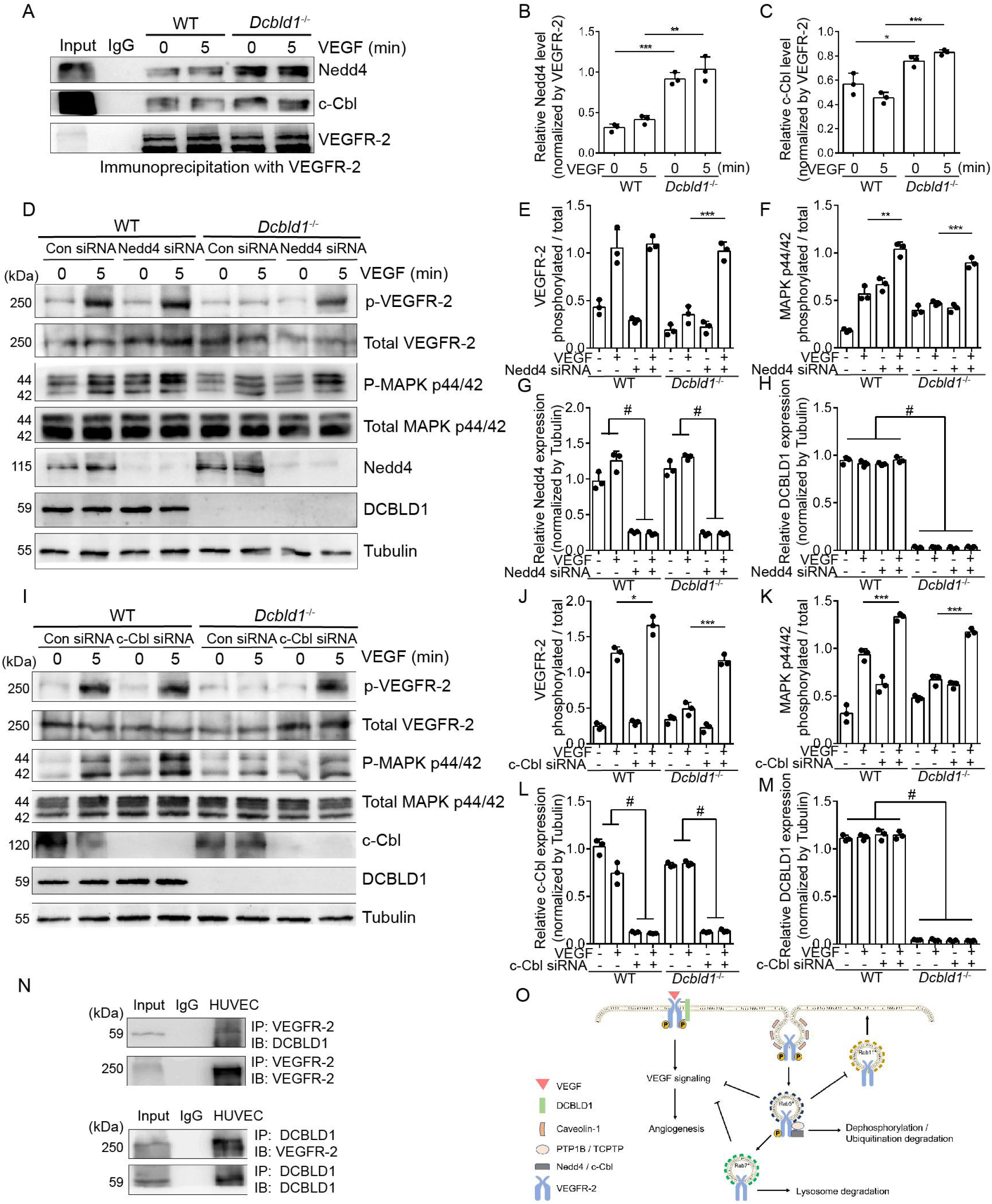
DCBLD1 deletion enhances ubiquitination of VEGFR-2 and influences VEGF signaling pathway in ECs. **(A-C)** Representative images of Western blot analysis after co-immunoprecipitation of E3 ubiquitin ligase, Nedd4 or c-Cbl, with VEGFR-2 in WT and *Dcbld1^-/-^* mPMVECs. *n=*3, \**p<*0.05, \*\**p<*0.01, \*\*\**p<*0.001. **(D-M)** Downregulated the expression of Nedd4 (D-H) or c-Cbl (I-M) proteins in WT and *Dcbld1^-/-^* mPMVECs by siRNA, analyzed the VEGFR-2 and p44/42 Thr^202^/Tyr^204^ MAPK phosphorylation induced by VEGF by Western blot, and performed quantitative analysis. *n=*3, \**p<*0.05, \*\**p<*0.01, \*\*\**p<*0.001, ^#^*p<*0.0001. **(N)** Verification of DCBLD1-VEGFR-2 protein interaction. The representative images of Western blot analysis after immunoprecipitation of VEGFR-2 and DCBLD1 in HUVECs. **(O)** The molecular mechanism of DCBLD1 on the dynamic regulation of VEGFR-2 recycling and degradation. DCBLD1 binds to VEGFR-2, promotes VEGF-induced activation of VEGFR-2 and downstream signaling pathways, and promotes angiogenesis. The absence of DCBLD1 increases the dephosphorylation and ubiquitination of VEGFR-2 after endocytosis and increases the degradation of VEGFR-2 through the lysosome pathway, thereby inhibiting VEGF signaling in ECs. The values are represented by mean±SEM. Statistical analyses were performed using two-way ANOVA, followed by Tukey’s multiple comparison test, compared to the control at the same time point.

To further address the mechanism of these tyrosine phosphatases or ubiquitinases in the effect of DCBLD1 deletion on VEGF signal transduction, we used siRNA to downregulate the expression of PTP1B, TCPTP, Nedd4, or c-Cbl in WT and *Dcbld1^-/-^* mPMVECs and quantified the status of VEGFR-2 andp44/42MAPK phosphorylation induced by VEGF. Western blot results showed that knockdown of PTP1B (Fig. 6D-H), TCPTP (Fig. 6I-M), Nedd4 (Fig. 7D-H), or c-Cbl (Fig. 7I-M) significantly enhanced VEGF-induced p44/42 MAPK phosphorylation and restored VEGFR-2 phosphorylation in *Dcbld1^-/-^* mPMVECs. These results showed that DCBLD1 could affect VEGF signal transduction by altering the dephosphorylation or ubiquitination degradation status of VEGFR-2 in ECs. Finally, we found that DCBLD1 can directly interact with VEGFR-2 in the co-immunoprecipitation experiment (Fig. 7N).

## Discussion

This study demonstrated that DCBLD1 combined with VEGFR-2 in ECs promotes VEGF-induced signal transduction and angiogenesis. The absence of DCBLD1 accelerates the Caveolin-1 mediated endocytosis of VEGFR-2, which increases retention in Rab5 vesicles, promotes dephosphorylation and ubiquitination of VEGFR-2, and promotes transport of VEGFR-2 to Rab7 positive vesicles. These effects increase the lysosomal degradation pathway of VEGFR-2 and reduce the recycling process of VEGFR-2 returning to the cell membrane through Rab11 positive vesicles, thereby weakening the VEGF signaling pathway of ECs and inhibiting angiogenesis (Fig. 7O). Therefore, we propose that DCBLD1 in ECs is a potential new regulator of development and adult angiogenesis.

VEGFR-2 is the main transmembrane receptor mediating VEGF function in ECs. After interacting with VEGF, the dimerization and subsequent phosphorylation of VEGFR-2 promote the endocytosis of the receptor and the downstream signaling pathway, thereby regulating the proliferation and migration of ECs, enhancing capillary permeability, and promoting the formation of peripheral blood vessels^30^. Previous studies have shown that DCBLD2 is a specific co-receptor of the same type of VEGF without intrinsic catalytic activity, which binds VEGF-A_165_ with VEGFR-2 to enhance VEGF signal transduction^31–33^. Full-length human sequences of DCBLD1 spans 715 amino acids. Each family member possesses a signal sequence, followed by CUB, LCCL, and coagulation factor V/VIII type-C (also Discoidin) domains^34,35^. A single-pass transmembrane region precedes the intracellular C-terminal scaffolding domain. The structure of the DCBLD domain closely resembles that of Neuropilins, which are transmembrane proteins that possess two CUB and discoidin domains and act as co-receptors for class 3 Semaphorins and growth factors in axon guidance and angiogenesis^10,36–38^. In this study, we demonstrated that downregulation of DCBLD1 inhibited VEGF-induced EC proliferation, migration, and angiogenesis. Studies have shown that VEGF-induced Akt and ERK activation supports EC survival and proliferation, whereas p38 phosphorylation mediates actin recombination and cell migration^21,30, 39–41^. We continued to track the effect of DCBLD1 on VEGF signaling pathway and found that the downregulation of DCBLD1 inhibited the phosphorylation of VEGFR-2, Akt, ERK, and p38 induced by VEGF.

Studies have shown that the changes in the biological behavior of ECs induced by abnormal VEGFR-2 activity are the cellular and molecular basis of many diseases^30,41^. The VEGFR-2 recycling process is a key step in regulating the activity of VEGFR-2, including the regulation of the dynamic balance of endocytosis and degradation of VEGFR-2^42–44^. The ligation of VEGFR-2 with VEGF on the cell surface induces entry into the cytoplasm mainly by the Clathrin-dependent pathway. However, when the pathway is inhibited, it can enter the cytoplasm through the Caveolar-dependent pathway^21^. Regardless the pathway by which VEGFR-2 enters the cells, the early endosomes contain Rab5 marker proteins. Rab5 recruits the intracellular adapter protein, APPL, to endosomes. APPL is a necessary adapter protein that activates downstream signaling molecules, ERK and Akt. Subsequently, EEA1 competes with APPL to bind Rab5 at the same site, and EEA1 gradually replaces APPL bound to early endosome marker, Rab5, which ultimately determines the subcellular localization of endosomes^21^. Under the influence of cytoskeletal proteins, such as Synectin, Kinesin-2, and Myosin-VI, endosomes or endosomes with Rab11 or Rab4 positive fusion, return to the cell membrane and complete the VEGFR-2 recycling process. Alternatively, the endosome marker, Rab5, is replaced by Rab7, and VEGFR-2 is transported to MVB and finally fused with lysosomes^22–26^. Based on these premises, our study addressed that the deletion of DCBLD1 promoted VEGFR-2 endocytosis by the caveolin-dependent pathway in HUVECs. VEGFR-2 entering the cytoplasm was mostly transported to Rab7-positive endosomes and, thus, transported to the MVB and degraded through the lysosome pathway. Few Rab11-positive endosomes fused to reduce the recycling process of VEGFR-2. The rapid degradation process of VEGFR-2 confirmed one of the internal mechanisms for the weakening of the VEGF signaling pathway after DCBLD1 deletion.

Among the regulators of VEGFR-2 activity, receptor ubiquitination is an important regulatory factor. Studies have shown that the ubiquitination of VEGFR-2 is mainly mediated by E3 ubiquitin ligase. E3 ubiquitin ligase family members, such as c-Cbl, Nedd4, and TRAF6, mediate the ubiquitination of VEGFR-2 binding to VEGF, promote the fusion and degradation of MVB and lysosome, and ultimately affect VEGF signaling^43,45–48^. The immunoprecipitation experiment on mPMVECs revealed that the deletion of DCBLD1 promoted the co-binding of VEGFR-2 with c-Cbl and Nedd4, indicating that the knockout of DCBLD1 accelerated the ubiquitination degradation of VEGFR-2. After downregulation of c-Cbl and Nedd4 in *Dcbld1^-/-^* mPMVECs, we restored the effect of loss of DCBLD1 in the reduction of VEGF signaling. This indicates that accelerating the ubiquitination degradation pathway of VEGFR-2 is one of the internal mechanisms for knocking out DCBLD1 to dampen the VEGF signaling pathway.

Interestingly, in the co-immunoprecipitation assay of mPMVECs, we found that deletion of DCBLD1 promoted the binding of VEGFR-2 to PTP1B and TCPTP, suggesting that deletion of DCBLD1 accelerated the dephosphorylation of VEGFR-2 and weakened the VEGF signaling pathway.

The co-immunoprecipitation of DCBLD1 and VEGFR-2 indicated that DCBLD1 and VEGFR-2 directly or indirectly interacted with part of the larger signal complex. According to a study, DCBLD2 (ESDN) or Neuropilin-1 binds to VEGFR-2 and regulates VEGFR-2 endocytosis^15,21^. Interestingly, VEGF-A_165_ forms complexes with VEGFR-2 and DCBLD2 (ESDN) or Neuropilin-1^15,49^, but the co-immunoprecipitation of DCBLD1 and VEGFR-2 does not require VEGF-A_165,_ which again indicated that more than one or two mechanisms of interaction exist. Unlike Neuropilin-1, DCBLD1 has a long cytoplasmic domain and several SH2 and SH3 binding sites, which may be involved in the interaction of other molecules. *In vivo*, unlike Neuropilin-1, *Neuropilin-1* gene deletion leads to embryonic lethality with defective vascular development^31,32^. However, *Dcbld1^-/-^* mice survive without any major vascular defects. In *Dcbld1^-/-^* mice, the inhibitory effects on development and adult angiogenesis were observed, indicating that the regulatory mechanism of DCBLD1 in the vascular system was different from that of Neuropilin-1.

Intimal hyperplasia is closely related to vascular remodeling disease, which is an important pathological basis of vascular wall thickening and luminal stenosis cardiovascular disease caused by endothelial injury, vascular smooth muscle cell proliferation, and extracellular matrix deposition^16,50^. This study has established that the expression of DCBLD1 plays an essential role in EC function. To further verify the effect of DCBLD1 expression on intimal hyperplasia, this study adopted the intimal hyperplasia model established by ligation of the common carotid artery of mice. H&E staining was used to measure intimal hyperplasia at different time points after ligation. The results showed that the degree of intimal hyperplasia in *Dcbld1^-/-^* mice was significantly higher than that in WT mice. This observation showed that DCBLD1 may affect intimal hyperplasia by affecting EC function. Tek-*Cre*^+/−^*Dcbld1*^flox/flox^ mice were used to further verify the effect of DCBLD1 in ECs on intimal hyperplasia. The results showed that, compared with the control mice, Tek-*Cre*^+/−^*Dcbld1*^flox/flox^ mice promoted vascular hyperplasia after vascular ligation, indicating that the EC function regulation controlled by DCBLD1 may play an important role in vascular remodeling.

In summary, our study demonstrated the essential role of endothelial DCBLD1 in regulating VEGF signal transduction and provided evidence that DCBLD1 promotes VEGF-induced angiogenesis by limiting the dephosphorylation, ubiquitination, and lysosome degradation of VEGFR-2. These findings provided new insights for endothelial DCBLD1 as a potential therapeutic target for ischemic cardiovascular diseases.

## Methods

### Animal experiments and ethics statement

All animal experiments were conducted according to the National Institutes of Health (NIH) Guide for the Care and Use of Laboratory Animals (NIH publication, 8th edition, 2011); they were approved by the Hebei Medical University Animal Care and Use Institutional Committee. All mice were fed at random in a 12-hour dark and 12-hour light cycle and a pathogen-free environment. The research protocol was submitted and approved by the Medical Ethics Committee of Hebei Medical University.

### Anaesthesia and euthanasia

For animal model and laser Doppler perfusion imaging, mice were anaesthetized with isoflurane (1-2%). At the end of the study, animals were euthanized by intraperitoneal injection of pentobarbital (80 mg/kg).

### Cell culture

Primary HUVECs were cultured in Dulbecco’s modified Eagle’s medium (DMEM), supplemented with 5% fetal bovine serum (FBS), 100 U/mL penicillin, and 100 μg/mL streptomycin. The cells were digested using trypsin containing EDTA and between 3–5 passages^42^. As described above, mPMVECs were isolated from WT and *Dcbld1*^-/-^ mice. PMVECs were purified after two rounds of immune selection using magnetic beads coated with anti-CD31 and anti-ICAM-2 antibodies (BD Bioscience, Franklin Lakes, NJ, USA). mPMVECs were cultured and maintained in a medium containing 5 % FBS and EC growth supplement (ECGS, Sigma-Aldrich, St. Louis, MO, USA). Unless otherwise stated, cells were used for experiments in early confluence.

### RNA interference assays

Three 25 bp siRNAs targeting human DCBLD1 mRNA were designed and synthesized by RiboBio:

stB0017177A: GUAUGACUGUUCCCAAAGA

stB0017177B: GUGGACACCUAGUGACUUA

stB0017177C: GGAACGAGCUAGCCAUUAU

siRNA was transiently transfected into HUVECs using Lipofectamine RNA iMAX transfection reagent (Thermo Fisher Scientific, Invitrogen, Carlsbad, CA, USA), according to the manufacturer’s protocol.

### Quantitative real-time PCR experiments

The total mRNAs were isolated using ABclonal kit and reverse transcribed. According to the manufacturer’s instructions, the cDNA was used for real-time PCR with three multiplex wells once with TaqMan gene assay kit (Thermo Fisher Scientific, Applied Biosystems, Carlsbad, CA, USA). The results were normalized to GAPDH. The TaqMan probes used in our study were DCBLD1 Hs 00543575_ml and GAPDH Hs 02758991_m1.

### Immunoprecipitation and Western blot experiments

After washing, the ECs were collected and lysed in RIPA buffer (150 mM NaCl, 50 mM Tris-HCl, pH 8.0, 1.0 % NP-40, 0.5 % sodium deoxycholate, and 0.1 % SDS) supplemented with protease inhibitors (Roche Applied Science, Manniheim, Germany) and phosphatase inhibitors (Sigma-Aldrich, St. Louis, MO, USA). Cell lysate concentration was determined using the DC Protein Assay Reagent (Bio-Rad Laboratories, Hercules, CA, USA). For immunoprecipitation, the cell lysate was incubated with 15 μL Dynabeads Protein A (Invitrogen, Thermo Fisher Scientific, Vilnius, LT, Norway) at 4°C for 1 h to pre-clear the non-specific protein binding. Thereafter, the cell lysate (500 μg) was immunoprecipitated with VEGFR-2 antibody at 4°C for 3 h and incubated with 20 μL Dynabeads overnight at 4°C. The same amount of cell lysates or immune precipitates was resolved by 10% SDS-PAGE and transferred to the PVDF membrane (Bio-Rad Laboratories, Hercules, CA, USA) by electrophoresis. The membrane was incubated with a primary antibody overnight at 4°C and with horseradish peroxidase (HRP)-conjugated secondary antibody (1:10000, Thermo Fisher Scientific, Waltham, MA, USA) thereafter for 1 h. The protein expression was visualized using Image Quant LAS4000 and quantified using ImageJ (NIH, Bethesda, MD, USA).

### Proliferation and migration assays

HUVEC proliferation was measured, as described in the product BrdU Cell proliferation Assay kit (Millipore, Temecula, CA, USA) and EdU Cell proliferation Assay kit (Beyotime, Shanghai, China). The migration of HUVECs was evaluated by the cell scratch assay. Cell migration rate represents the area of cells migrating to the wounded area in each field of vision. Concurrently, cell migration was evaluated by an improved Borden chamber assay. HUVEC suspension (1×10^5^ cells) was added in the upper chamber of Transwell (3422, Corning Incorporated, Corning, NY, USA), and 0.5% FBS and VEGF-A_165_ (100 ng/mL) were added to the lower chamber. After migration in a 37°C incubator for 12 h, the transwell upper chamber was washed with PBS, and the cells in the upper chamber were wiped with cotton swabs. The cells at the bottom of the Transwell were fixed with 4% paraformaldehyde (PFA) and stained with 0.1% crystal violet. Leica phase contrast light microscope was used to produce images of eight random microscopic fields of each Transwell, and ImageJ was used to calculate the number of migrated cells.

### *In vitro* tube formation assay

The cells were starved overnight in 0.5% FBS. Thereafter, they were treated with trypsin and added to a 48-well plate with pre-prepared matrix (Corning Incorporated, Bedford, MA, USA) at a density of 60,000 cells per well. The ECs were cultured in the conditioned medium containing 0.5% FBS or the medium containing 0.5% FBS and VEGF-A_165_ (100 ng/mL) for 6 h. The cumulative tube length in five random fields was measured under a phase contrast microscope to quantify the *in vitro* angiogenetic ability of these ECs. The average cumulative total length of each hole represented an experimental point, and ImageJ was used to evaluate tubule branch number.

### Immunofluorescence microscopy

For tissue immunofluorescence staining, fresh frozen OCT tissue sections (5 μm thickness) were fixed in precooled acetone (–20 °C) for 5 min, washed three times in PBS, and incubated overnight at 4°C with primary antibodies against DCBLD1 (1:30, Atlas antibody, Bromma, Sweden) and CD31 (1:100, BD Bioscience, Franklin Lakes, NJ, USA). Thereafter, the samples were incubated with Alexa Fluor 488 or Alexa Fluor 594-conjugated secondary antibodies (1:100, Thermo Fisher Scientific, Waltham, MA, USA) for 1 h. The sections were mounted using ProLong Gold antifade medium containing DAPI (Thermo Fisher Scientific, Waltham, MA, USA). Leica SP2 confocal microscopy (Leitz, Wetzlar, Germany) was used to examine fluorescence images.

For cell immunofluorescence staining, HUVECs on the coverslip were rinsed twice in cold PBS, fixed in freshly prepared 4% PFA for 30 min at room temperature, rinsed three times in PBS (containing 0.1% Tween-20), sealed in PBS (containing 0.05 % TritonX-100 and 5% BSA) for 30 min, rinsed three times in PBS (containing 0.1 % Tween-20), incubated with primary antibody overnight at 4°C, and rinsed three times in PBS (containing 0.1 % Tween-20). Thereafter, cells were incubated with Alexa Fluor 488 or Alexa Fluor 594-coupled secondary antibodies for 1 h. Slides were then washed with PBS three times, mounted, and examined, as described previously.

### Generation of DCBLD1 genetically engineered mouse model (GEMM) and genotyping

Gene knockout *Dcbld1^-/-^* mice were developed by the Animal Center of Hebei Medical University. The EC-specific conditional DCBLD1 knockout mice were developed by crossing *Dcbld1*^flox/flox^ and Tek-Cre transgenic mice (generated by Biocytogen Pharmaceuticals, Beijing, China). Mice at 6–8 weeks of age were crossed at a female-to-male ratio of 2:1. The genomic DNAs extracted from the ears and genotypes were identified by PCR to confirm that Tek-Cre^+/−^*Dcbld1*^flox/flox^ mice and their controls, *Dcbld1*^flox/flox^ mice, had been successfully developed.

Primers for genotyping the whole genome of *Dcbld1^-/-^*, Tek-Cre^+/−^*Dcbld1*^flox/flox^, and control littermate *Dcbld1*^flox/flox^ mice are as follows: *Dcbld1* primers designed by deleting sequences from DCBLD1. PCR product size: 225 bp.

Forward primer: 5’-TAGGCCAAAAGTCCCTATTGC-3’

Reverse primer: 5’-TGCTGGAGTTACCAATTAGTCG -3’

*Dcbld1* 5’loxP primers. PCR product size: 468bp, wild type: 356 bp.

Forward primer: 5’-GCCCTGGCTATCCTGGAACTTTGTA-3’

Reverse primer: 5’-AGAGAACATACTCTGGGATCCTCCT-3’

*Dcbld1* 3’loxP primers. PCR product size: 484bp, wild type: 377 bp.

Forward primer: 5’-CCTGGTTGGTATTTGGAAATAAGGTGTA-3’

Reverse primer: 5’-TCTAGACAGAAATTGCCCGCTGGAG-3’

Tek-*Cre*^+/−^primers. PCR product size: positive control 324 bp, Cre 100 bp.

Tek-*Cre* forward primer: 5’-GCGGTCTGGCAGTAAAAACTATC-3’

Tek-*Cre* reverse primer: 5’-GTGAAACAGCATTGCTGTCACTT-3’

Tek-*Cre* positive control

forward primer: 5’-CTAGGCCACAGAATTGAAAGATCT-3’

Tek-*Cre* positive control

reverse primer: 5’-TAGGTGGAAATTCTAGCATCATC-3’

### Common carotid artery ligation model

WT, *Dcbld1*^-/-^, Tek-Cre^+/−^*Dcbld1*^flox/flox^, and their control littermate *Dcbld1*^flox/flox^ mice were randomly divided into five groups (control, 3, 7, 14, and 21 days after ligation). An intimal hyperplasia model was established by ligating the left common carotid artery. The surgery was performed under isoflurane inhalation anesthesia with the aid of a microscope. In the sham control group (sham surgery), blood vessels were separated without ligation. On the 3rd, 7th, 14^th^, and 21st days after ligation, the left common carotid artery was collected after perfusion with normal saline, placed in OCT embedding agent (Tissue Tek, Torrance, CA, USA), and quickly frozen in liquid nitrogen for H&E staining or immunofluorescence staining.

### Hind limb ischemia model and laser Doppler perfusion imaging

Male mice (6–8 weeks old) were anesthetized with isoflurane for unilateral hind limb surgery. We performed ligation and segmental resection of the left femoral vessels, followed by physiological and histological analyses^51^. Briefly, the left femoral artery was exposed and ligated with 6-0 sutures at the proximal and distal ends. The blood vessels between the ligation lines were removed without damaging the femoral nerve.

We used a laser Doppler flow analyzer (Peri Scan PIM system) to analyze the lower limb blood flow of mice before and immediately after surgery and on the 3rd, 7^th^, and 14th days after surgery. Additionally, the ischemic (left)/non-ischemic (right) lower limb blood flow ratio was measured. Before the measurement, the mice were anesthetized and placed on a heating plate at 37°C for 10 min to minimize the temperature variations. On the 14^th^ day post-surgery, the mice were euthanized by injecting pentobarbital (80 mg/kg)^52^, and the gastrocnemius muscle tissue was obtained and placed in OCT embedding agent. The frozen sections (5 μm thickness) were used for immunofluorescence analysis.

### Retinal angiogenesis assay

The postnatal eyes of mice on the fifth day of birth were fixed in 4% PFA for 30 min and washed. The retina was dissected in PBS and then permeabilized overnight in PBS containing 1% BSA and 0.5% TritonX-100. The transparent retina was incubated with the FITC-conjugated Isolectin-B4 (20 μg/mL, Enzo Life Science, Farmingdale, NY, USA). Using the Biologic CMM Analyzer software (French College), the total length and number of branches of co-immunoreactive blood vessels throughout the retina were quantified on a composite magnification image.

### Isolation of plasma-membrane subfraction

To explore the role of DCBLD1 on VEGFR-2 localization, HUVECs were starved in DMEM for 24 h and then stimulated with VEGF-A_165_ (10 ng/mL) for the corresponding time to collect cellular fractions. According to manufacturer’s instructions, cytoplasmic and cell membrane components were prepared using cell fraction isolation kits (Invent is Bio, Shanghai, China).

### Co-immunoprecipitation (Co-IP) and mass-spectrum (MS) assays

The Total protein was extracted from WT or *Dcbld1*^-/-^ mPMVECs pretreated with VEGF-A_164_ using modified RIPA buffer (150mM NaCl, 50mM Tris-HCl, pH8.0, 1.0 % NP-40, 0.5 % sodium deoxycholate and 0.1 % SDS), and co-immunoprecipitation with VEGFR-2 antibody, SDS-PAGE was performed. The IP products (immunoprecipitated with VEGFR-2) were analyzed by MS. MS was conducted by equipments of Q Exactive (Thermo Fisher) and Easy-nLC 1000 (Thermo Fisher). The data of MS were analyzed by MASCOT Software. Select differentially expressed proteins from parts with Fold Change values greater than 2 or less than 0.5. To explore biological processes associated with DCBLD1-interacting proteins, enrichment analysis in the GO domain BP, CC and MF were performed.

### Statistical analysis

The data were expressed as mean±SEM. Prism 6 (GraphPad Software, San Diego, CA) was used for statistical analyses. Data were compared between cell group and animal group using the two-tailed unpaired Student *t* test or two-way ANOVA, followed by Tukey’s multiple comparison test. In statistical analysis, *n* represents the number of independent biological samples or mice. *P<*0.05 were considered statistically significant.

## Acknowledgments

This work was supported by the National Natural Science Foundation of China (31571176 and 31771268 to L.N, 81600576 to H.N), Natural Science Foundation of Hebei Province of China (2016206226 to L.N), Precision medicine joint fund of Hebei Province of China (2020206432 to L.N, 2021206393 to L.N).

## ETHICAL STATEMENTS

All the gene modify and control littermate mice were raised in the Laboratory Animal Centre of Hebei Medical University. All animal experiments were performed following the guidelines for laboratory animal manipulation approved by the Hebei Laboratory Animals Monitoring Institute and under the inspection of the Animal Welfare and Ethics Review Board of Hebei Medical University.

## AUTHOR CONTRIBUTIONS

**Qi Feng:** Conceptualization, Methodology, Formal analysis, Writing-Original draft preparation **Lingling Guo:** Conceptualization, Methodology, Formal analysis, Data curation **Chao Yu:** Conceptualization, Methodology, Formal analysis, Data curation **Xiaoning Liu:** Conceptualization, Methodology, Formal analysis **Yanling Lin:** Methodology, Formal analysis **Chenyang Li:** Methodology, Validation **Wenjun Zhang:** Methodology, Validation **Yanhong Zong:** Methodology, Formal analysis **Weiwei Yang:** Methodology **Yuehua Ma:** Methodology **Runtao Wang:** Methodology **Lijing Li:** Methodology **Yunli Pei:** Methodology, Validation **Huifang Wang:** Validation, Software **Demin Liu:** Resources, Software **Mei Han:** Methodology, Resources **Honglin Niu:** Resources, Funding acquisition, Project administration **Lei Nie:** Investigation, Supervision, Funding acquisition, Writing-Review & Editing. All authors have read and agreed to the published version of the manuscript.

## DATA AVAILABILITY STATMENT

The publisher is not responsible for the content or functionality of any supporting information supplied by the authors. The data that support the findings of this study are available on request from the corresponding author.

---

**Supplemental Figure 1.**
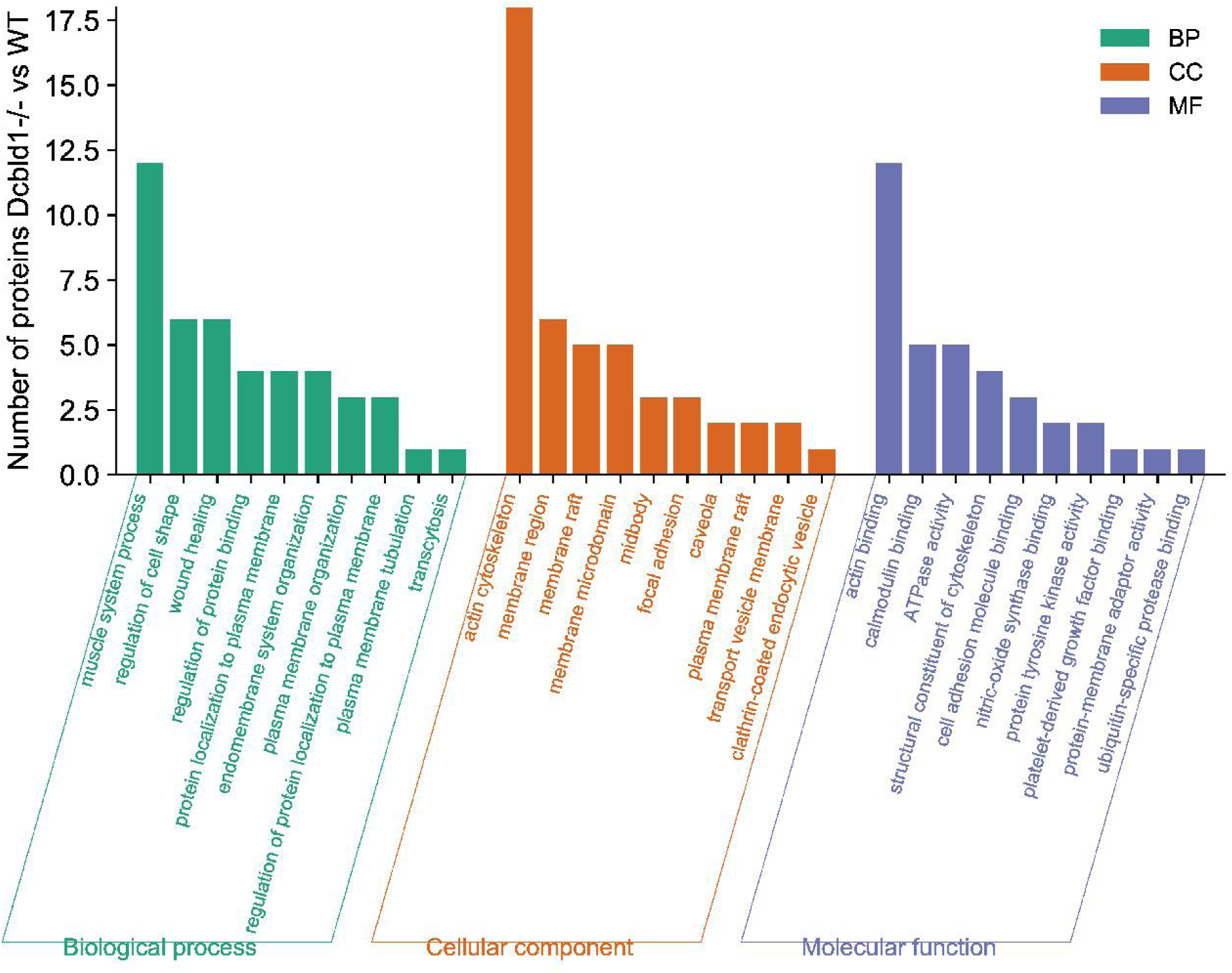
Gene Ontology (GO) enrichment analysis. An overview of top 10 significantly enriched terms in three categories: biological process (BP), cellular component (CC), and molecular function (MF). Number of proteins involved a process is shown in the Y-axis. The cut-off of *P*-value was set to 0.05.

**Supplemental Figure 2.**
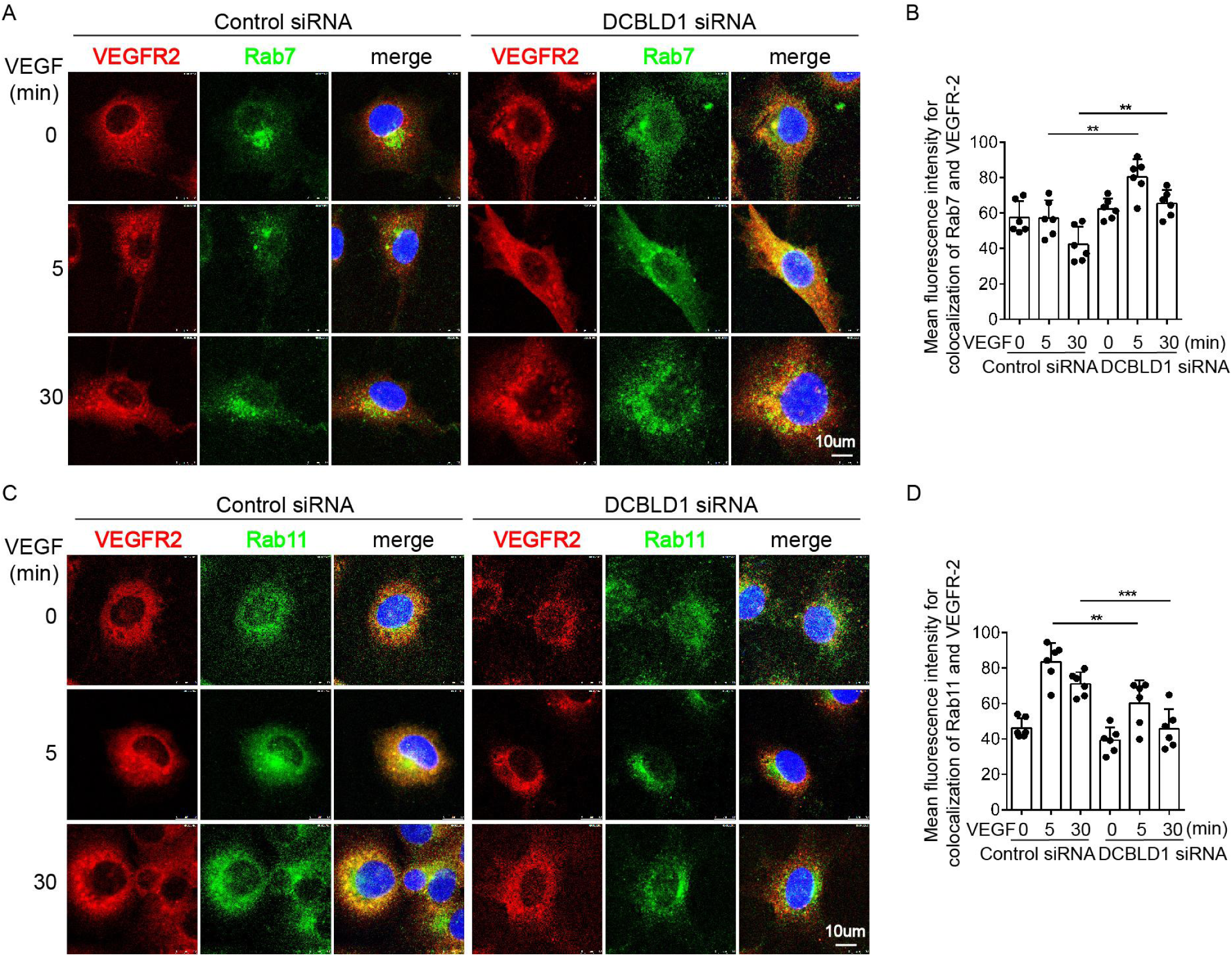
Downregulation of DCBLD1 affects the binding of VEGFR-2 to Rab7 and Rab11 in HUVECs. **(A, B)** After DCBLD1 siRNA transient transfection, HUVECs were treated with VEGF (10 ng/mL) at different time points. VEGFR-2 (red) and Rab7 (green) representative images of immunofluorescence staining and the nucleus with DAPI (blue) staining. Bar=10 μm. *n=*6,\*\**P<*0.01.**(C, D)** After DCBLD1 siRNA transient transfection, HUVECs were treated with VEGF (10 ng/mL) at different time points. VEGFR-2 (red) and Rab11 (green) representative images of immunofluorescence staining and the nucleus with DAPI (blue) staining. Bar=10 μm. *n=*6, \*\**P<*0.01, \*\*\**P<*0.001.The values are represented by mean± SEM. Statistical analyses were performed using two-way ANOVA, followed by Tukey’s multiple comparison test, compared to the control at the same time point.

**Supplemental Table 1.**
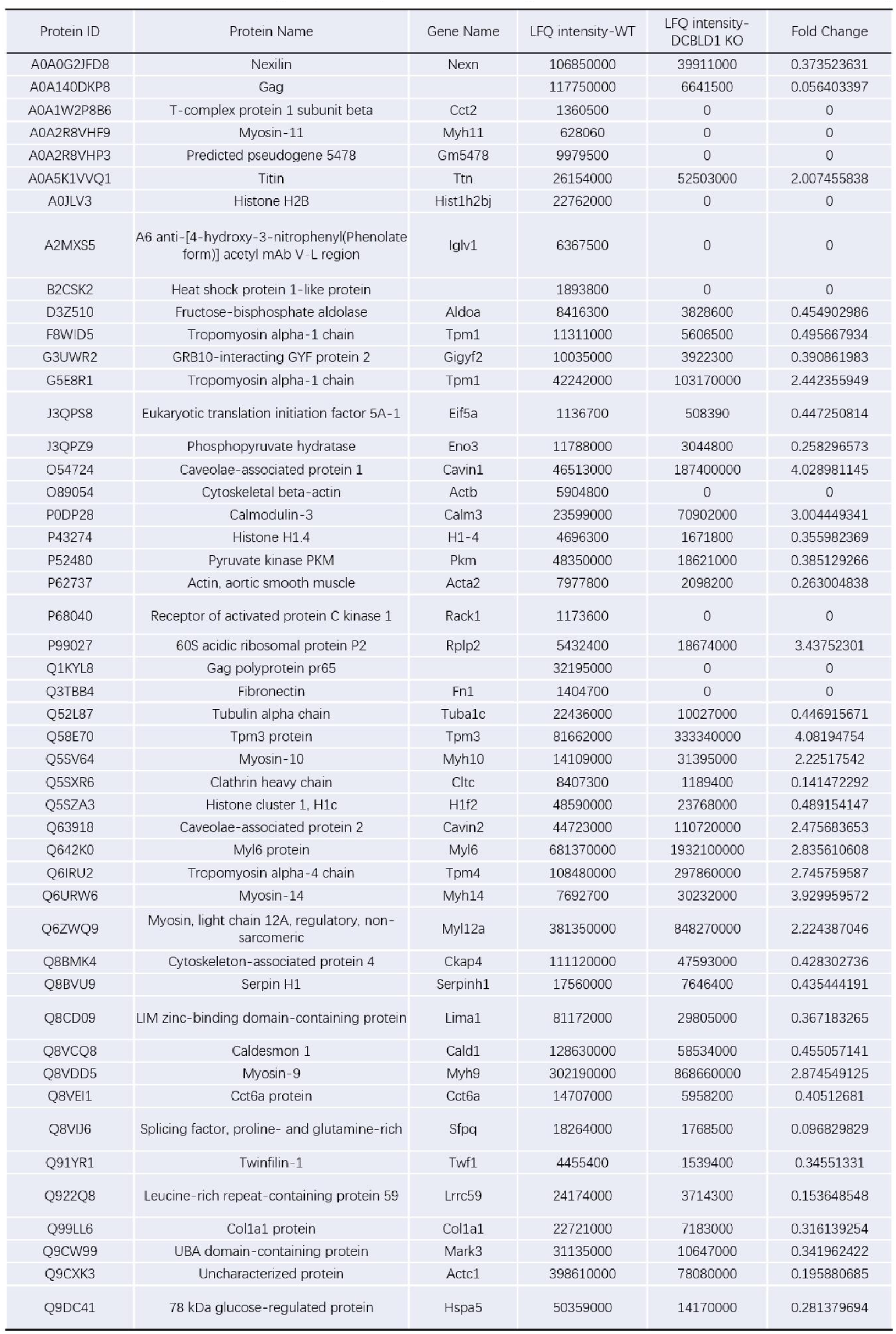
MS identified differentially expressed proteins interacting with VEGFR-2 in WT or *Dcbld1*^-/-^mPMVECs specific bands. The proteins co-binding with VEGFR-2 in WT or *Dcbld1*^-/-^mPMVECs were detected by MS. The differential proteins were selected from the part with Fold Change value >2 or <0.5.

